# Human Auditory Ossicles as an Alternative Optimal Source of Ancient DNA

**DOI:** 10.1101/654749

**Authors:** Kendra Sirak, Daniel Fernandes, Olivia Cheronet, Eadaoin Harney, Matthew Mah, Swapan Mallick, Nadin Rohland, Nicole Adamski, Nasreen Broomandkhoshbacht, Kimberly Callan, Francesca Candilio, Ann Marie Lawson, Kirsten Mandl, Jonas Oppenheimer, Kristin Stewardson, Fatma Zalzala, Alexandra Anders, Juraj Bartík, Alfredo Coppa, Dashtseveg Tumen, Sándor Évinger, Zdeněk Farkaš, Tamás Hajdu, Jamsranjav Bayarsaikhan, Lauren McIntyre, Vyacheslav Moiseyev, Ildikó Pap, Michael Pietrusewsky, Pál Raczky, Alena Šefčáková, Andrei Soficaru, Tamás Szeniczey, Béla Miklós Szőke, Tumurbaatar Tuvshinjargal, Dennis Van Gerven, Sergey Vasilyev, Lynne Bell, David Reich, Ron Pinhasi

## Abstract

DNA recovery from ancient human remains has revolutionized our ability to reconstruct the genetic landscape of the past. Ancient DNA research has benefited from the identification of skeletal elements, such as the cochlear part of the osseous inner ear, that provide optimal contexts for DNA preservation; however, the rich genetic information obtained from the cochlea must be counterbalanced against the loss of valuable morphological information caused by its sampling. Motivated by similarities in developmental processes and histological properties between the cochlea and auditory ossicles, we evaluated the efficacy of ossicles as an alternative source of ancient DNA. We demonstrate that ossicles perform comparably to the cochlea in terms of DNA recovery, finding no substantial reduction in data quality, quantity, or authenticity across a range of preservation conditions. Ossicles can be sampled from intact skulls or disarticulated petrous bones without damage to surrounding bone, and we argue that, when available, they should be selected over the cochlea to reduce damage to skeletal integrity. These results identify a second optimal skeletal element for ancient DNA analysis and add to a growing toolkit of sampling methods that help to better preserve skeletal remains for future research while maximizing the likelihood that ancient DNA analysis will produce useable results.

## INTRODUCTION

Ancient DNA has become an important tool for addressing key questions about human evolutionary and demographic history. Its rapid growth over the last decade has been driven largely by advances in isolating (Dabney et al. 2013; Rohland et al. 2018), preparing (Gansauge et al. 2017; Rohland et al. 2015), enriching (Fu et al. 2013, 2015; Haak et al. 2015; Mathieson et al. 2015), sequencing (Margulies et al. 2005), and analyzing (Briggs et al. 2007; Briggs et al. 2010; Ginolhac et al. 2011; Skoglund et al. 2014) small quantities of degraded DNA. While these methodological advances have contributed to an improvement in the quality and quantity of paleogenomic data obtained from ancient human remains, all ancient DNA research fundamentally depends upon access to biological material that has sufficient biomolecular preservation.

Skeletal tissue (i.e., bone or teeth) is the preferred biological material for human ancient DNA analysis due to its ability to resist *post-mortem* degradation better than other types of tissues, including skin and hair (Lindahl 1993; Smith et al. 2001, 2003; Collins et al. 2002). Recent research has shown that not all bone elements are equally effective in preserving DNA, however, and has identified the bone encapsulating the cochlea within the petrous pyramid of the temporal bone (referred to henceforth as the ‘cochlea’) (Gamba et al. 2014; Pinhasi et al. 2015), as well as the cementum layer in teeth roots (Damgaard et al. 2015; Hansen et al. 2017) as especially DNA-rich parts of the skeleton. The use of these skeletal elements that act as repositories for the long-term survival of DNA has proven to be particularly important for the analysis of biological samples recovered from regions where high temperatures and/or humidity increase the rate of molecular degradation and result in low concentrations of damaged DNA with reduced molecular complexity (e.g., Broushaki et al. 2016; Lazaridis et al. 2016; Schuenemann et al. 2017; Skoglund et al. 2017; Fregel et al. 2018; Harney et al. 2018; van de Loosdrecht et al. 2018).

While use of the cochlea has contributed to the application of ancient DNA research to a growing range of geographic and temporal contexts, it is important to balance analytical goals with the irreparable damage to human skeletal remains that results from destructive analyses (Prendergast and Sawchuk 2018; Sirak and Sedig *in press*). Ancient DNA is one of several such analyses that are now widely used in archaeology (others include radiocarbon dating and stable isotope analysis) (Hublin et al. 2008; Mays et al. 2013; Makarewicz et al. 2017; Pinhasi et al. 2019). To minimize damage to intact skulls from ancient DNA sampling while still accessing the rich genetic data in the cochlea, we developed a “Cranial Base Drilling” method to minimize damage to surrounding bone areas when a skull is intact (Sirak et al. 2017). However, even this method involves destructive sampling. Recent work has highlighted the fact that morphological analysis of the inner ear part of the petrous pyramid (including the cochlea) can reveal population relationships and thus harbors some information about population history (e.g., Spoor et al. 2003; Ponce de León et al. 2018). While genetic comparisons of samples involve analysis of tens of thousands of independent markers (single nucleotide polymorphisms, or SNPs) which provide far higher statistical resolution than can be obtained by study of the smaller number of data points that can be extracted from morphological analysis, not all cochlear bone yields sufficient amounts of ancient DNA. The fact that there is morphological information in the petrous pyramid that will be destroyed through sampling of ancient DNA highlights the importance of being a careful steward of these elements.

As part of a search for alternative optimal sources for ancient DNA that can be used in place of the cochlea, we noted that auditory ossicles have similar developmental processes and histological properties as the osseous inner ear. We therefore tested whether the ossicles – the smallest bones in the human body – might serve as alternative optimal substrates for ancient DNA analysis.

### Ossicle development and histology

The mechanism by which cochlear bone preserves endogenous DNA better than other skeletal elements or other regions of the same petrous pyramid is not well understood; however, it is likely related to the fact that human petrous bones are unique in being characterized by a near-absence of growth or remodeling following the completion of ossification by approximately 24 weeks *in utero* (Sølvsten Sørensen et al. 1992; Frisch et al. 1998; Hernandez et al. 2004). The inhibition of bone remodeling leads to the presence of a larger number of mineralized osteocytes that reside in lacunae within the bone tissue (Hernandez et al. 2004; Bell et al. 2008; Busse et al. 2010; Rask-Andersen et al. 2012). One hypothesis (Pinhasi et al. 2019) is that ‘microniches’ created in the bone tissue by the maintenance of mineralized osteocytes, combined with the protected location of the cochlea, may act as repositories that encourage the long-term preservation of DNA (Bell et al. 2008; Kontopoulos et al. 2019). Ossicles are similar to the cochlea in this respect (see below), and we therefore hypothesized that they might also preserve high amounts of endogenous DNA.

In humans, the middle ear (the region of the ear located medial to the eardrum and lateral to the oval window of the inner ear) is enclosed within the temporal bone and contains the three auditory ossicles: the malleus, incus, and stapes (Figure 1). The ossicles effectively allow humans to hear by transmitting sound-induced mechanical vibrations from the outer to the inner ear. Though the ossicles do not experience high-strain biomechanical loading, they are subject to unique vibrational patterns that impact their development and characteristics over the course of an individual’s lifespan (Rolvien et al. 2018). In contrast to the majority of the human skeleton, but similar to the cochlea, the auditory ossicles present with their final size and morphology at birth following the onset of the ossification of between 16 and 18 weeks *in utero* and the completion of ossification around 24 weeks gestational age (Marotti et al. 1998; Yokoyama et al. 1999; Cunningham et al. 2000; Duboeuf et al. 2015; Richard et al. 2017). The ossicles and cochlea appear to follow the same developmental pattern of rapidly increasing bone volume through cortical thickening and densification, along with mineralization of the bony matrix (Richard et al. 2017).

**Figure 1:**
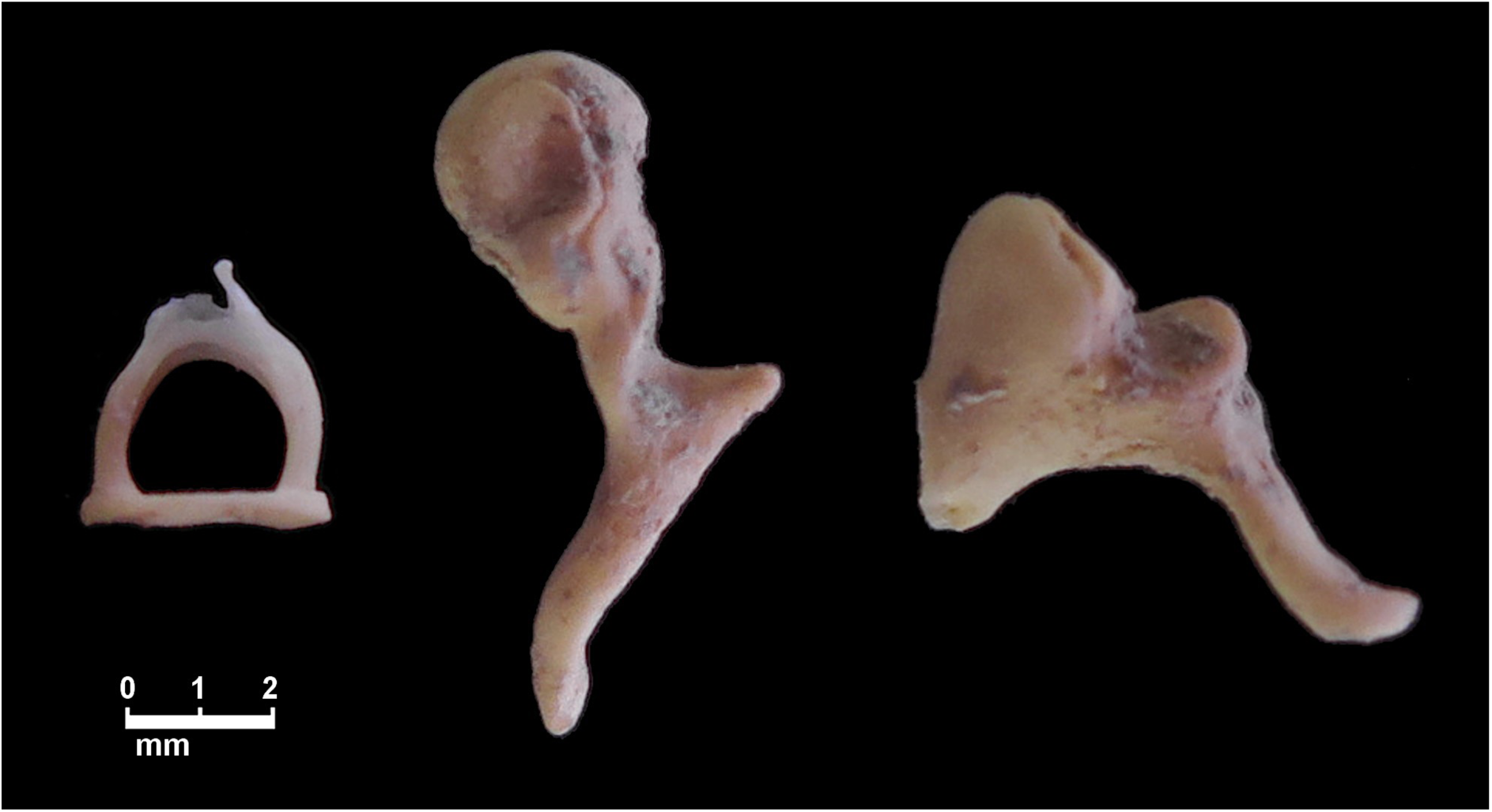
The three auditory ossicles. From left to right, the stapes, malleus, and incus.

Like the cochlea, ossicular bone tissue is rapidly modeled around the time of birth; although it may undergo further postnatal maturation, there are no signs of bone remodeling observed above the age of one year (Richard et al. 2017; Rolvien et al. 2018). The inhibition of bone remodeling of the auditory ossicles is evident from features such as the presence of a dense meshwork of collagenous fibers organized in an interlacing woven pattern, a smooth fibrous appearance, and limited vascular channels and viable osteocytes (Marotti et al. 1998; Chen et al. 2008). As in the case of the cochlea and in contrast to other skeletal elements, mineralized osteocytes appear to accumulate in the ossicles throughout an individual’s life without resulting in increased bone absorption (Marotti et al. 1998; Kanzaki et al. 2006; Rolvien et al. 2018), likely conserving the overall architecture of the ossicles in order to maintain optimal sound transmission (Kanzaki et al. 2006; Rolvien et al. 2018). While the consequences of inhibited bone remodeling and the accumulation of mineralized osteocytes have only been previously studied from a clinical perspective, we hypothesized that these features might contribute to optimized DNA preservation similar to that in the cochlea by creating the ‘microniches’ that enable long-term DNA survival (Bell et al. 2008).

### Use of ossicles in ancient DNA research

Due to their small size and tendency to become dislodged from the skull, ossicles are only seldom recovered during excavation and are easily lost in collections excavated decades ago. While ossicles are not recovered for every burial in every context, we have empirically found that these bones may remain lodged within the middle ear of intact skulls or can be identified in the vicinity of a burial during excavation (Qvist et al. 2000). Given the value of the ossicles as a substrate for ancient DNA analysis, demonstrated in this study, we hope that more archaeologists and anthropologists and museum curators will focus on preserving these elements.

It is important to recognize that ossicles, just like the cochlea, are morphologically informative. Indeed, there is a growing body of literature examining the comparative morphology and pathology of the ossicles (e.g., Rak and Clarke 1979; Arensburg et al. 1981, 2005; Siori et al. 1995; Spoor et al. 2003; Crevecoeur et al. 2007; Quam and Rak 2008; Quam et al. 2013a, 2013b; Stoessel et al. 2016). While differences in metric and non-metric features of the auditory ossicles may be taxonomically informative for comparisons across the genus *Homo* (e.g., Heim 1982; Spoor et al. 2003; Quam and Rak 2008; though see Arensburg et al. 1981), it is unclear whether phylogenetic and population relationship information can be retrieved from the auditory ossicles. In cases where ossicle morphology may be a subject of future research, we encourage that anthropological study (including description, measurement, and evaluation of any apparent pathologies) and surface or micro-CT scanning to collect metric and morphological information prior to ancient DNA analysis. Any ossicles that exhibit visible pathologies should be avoided.

Though some anthropological attention has been given to the ossicles, we are not aware of previous genetic analyses of these bones. Only a single study has attempted to analyze DNA from the ossicles, collecting the ossicles during medical autopsies of recently-deceased individuals and determining them to be a reliable DNA source from bodies ranging from freshly deceased to highly putrefied (Schwark et al. 2015).

## RESULTS

We carried out pilot work to assess if the quality and quantity of ancient DNA data recovered from the ossicles was approximately similar to that recovered from the cochlea (described in Supplemental Material). The results of this pilot work (Supplementary Table 1) suggested that ossicles perform comparably to the cochlea in metrics such as amount of endogenous human DNA recovered and frequency of damage at the terminal nucleotide of the DNA molecule (a commonly used measure of ancient DNA authenticity). Based on these results we selected 10 ossicles from archaeological samples from a wide range of geographic locations with varying climates and dated to between ∼6500–1720 years before present (yBP) (Table 1, with detailed sample information in Supplementary Table 2). To be included in this study, each specimen was required to have at least one ossicle as well as the cochlea of the petrous bone available for comparative analysis. Whenever possible, a petrous bone that had an antimere was chosen (Prendergast and Sawchuk 2018); we did not sample the antimeres in order to preserve them for future analyses.

**Table 1:** Sample information and summary of sequencing results.

| ID (I-ID) | Site (Country) | Date | Initial Material | Shotgun Endogenous % (hg19) | 1240K SNPs | 1240K Coverage | 1240K Terminal Base Deamination | mtDNA Consensus Match [95CI] | 1240K ANGSD Contamination Mean [Z-score] | % Unique Reads Expected at 500K On-Target Sequences |
| --- | --- | --- | --- | --- | --- | --- | --- | --- | --- | --- |
| 614 | Katelai (Pakistan) | 1000-800 BCE | Powder 56 mg | 7.527% | 622982 | 0.541 | 0.139 | 0.996 [0.990-0.999] | 0.008 [2.897] | 67.8 |
|  |  |  | Ossicle (2: I, M) | 30.550% | 776019 | 0.674 | 0.131 | 0.994 [0.986-0.998] | 0.005 [2.593] | 83.2 |
| 644 | Geoksiur (Turkmenistan) | 5000-2000 BCE | Powder 47 mg | 3.270% | 391574 | 0.340 | 0.239 | 0.994 [0.984-0.998] | N/A (Female) | 50.4 |
|  |  |  | Ossicle (2: I, M) | 0.567% | 177819 | 0.155 | 0.225 | 0.989 [0.974-0.996] | N/A (Female) | 31.5 |
| 648 | Fofonovo (Russia) | 6000-3000 BCE | Powder 47 mg | 70.145% | 829510 | 0.721 | 0.044 | 0.999 [0.995-1.000] | 0.002 [2.053] | 98.9 |
|  |  |  | Ossicle (2: I, S) | 72.675% | 858204 | 0.746 | 0.045 | 0.991 [0.983-0.996] | 0.002 [3.254] | 98.8 |
| 818 | Bayankhongor, Ulziit sum, Maanit uul (Mongolia) | 2500-400 BCE | Powder 55 mg | 56.484% | 855679 | 0.744 | 0.139 | 0.995 [0.987-0.998] | 0.005 [5.841] | 98.7 |
|  |  |  | Ossicle (1: I) | 71.801% | 883674 | 0.768 | 0.106 | 0.965 [0.953-0.974] | 0.040 [20.256] | 98.2 |
| 1257 | Peugeot Garage (Great Britain) | 43-410 CE | Powder 49 mg | 46.785% | 843283 | 0.733 | 0.134 | 0.981 [0.972-0.988] | N/A (Female) | 97.7 |
|  |  |  | Ossicle (2: I, M) | 55.833% | 829630 | 0.721 | 0.128 | 0.971 [0.960-0.980] | N/A (Female) | 94.6 |
| BCB 26 | Ban Chiang (Thailand) | 900-300 BCE | Powder 48 mg | 0.010% | 266 | 0.000 | 0.064 | - | N/A (Female) | - |
|  |  |  | Ossicle (3: I, M, S) | 0.068% | 12438 | 0.008 | 0.013 | 0.940 [0.877-0.972] | N/A (Female) | - |
| BLAT 33 | Blatné (Slovakia) | 2200-2000 BCE | Powder 50 mg | 63.000% | 770152 | 3.909 | 0.091 | 0.996 [0.990-0.999] | 0.006 [6.180] | 98.6 |
|  |  |  | Ossicle (2: I, M) | 68.250% | 725438 | 3.914 | 0.088 | 0.998 [0.993-1.000] | 0.005 [5.184] | 99.1 |
| GLAV 14 | Glăvănești (Romania) | 3500-3000 BCE | Powder 47 mg | 73.877% | 779326 | 3.727 | 0.078 | 0.995 [0.989-0.998] | 0.005 [4.856] | 97.9 |
|  |  |  | Ossicle (2: I, M) | 59.397% | 749401 | 3.343 | 0.068 | 0.991 [0.982-0.995] | 0.004 [4.574] | 96/8 |
| MKDH 13H | Al-Makhdarah (Yemen) | 1190-800 BCE | Powder 53 mg | 0.002% | 253 | 0.000 | 0.014 | - | - | - |
|  |  |  | Ossicle (2: I, M) | 0.001% | 531 | 0.000 | 0.025 | - | - | - |
| KHRB UA | Kharibat al-Ahjur (Yemen) | 1000 BCE | Powder 53 mg | 0.004% | 115 | 0.000 | 0.024 | - | - | - |
|  |  |  | Ossicle (2: I, M) | 0.007% | 1244 | 0.001 | 0.098 | - | - | - |

A summary of sequencing results for the 10 individuals reported in this paper is presented in Table 1 and Figure 2; for more detailed information, see Supplementary Table 2.

**Figure 2:**
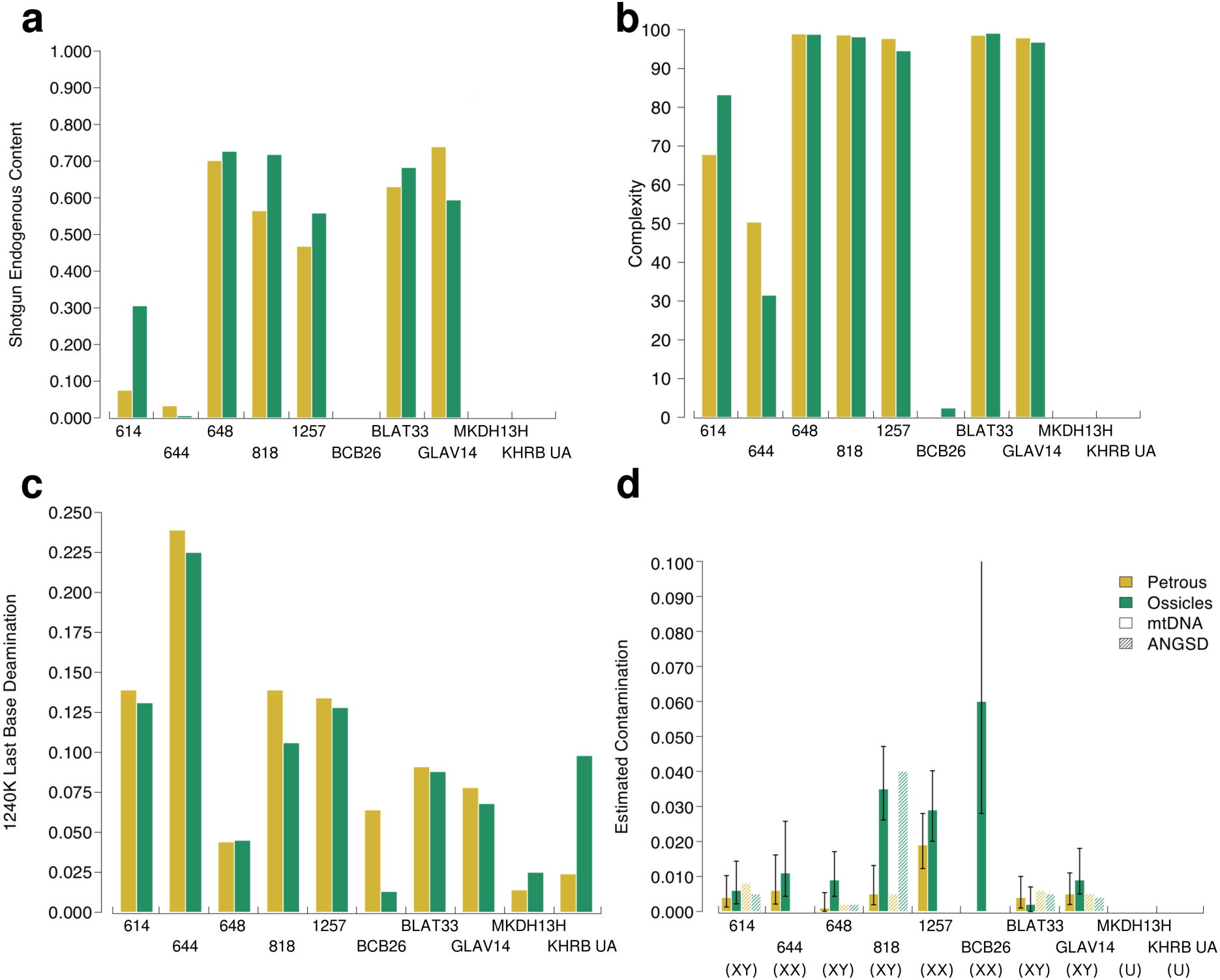
Comparative results between cochlea (yellow) and ossicle (green) samples from the same individuals. **Panel a.** Endogenous shotgun DNA ratios of the total reads. **Panel b.** Complexity as percentage of unique reads expected from 500,000 reads hitting targets. **Panel c.** Deamination frequencies on the terminal bases of the 1240K capture sequences. **Panel d.** Contamination estimates calculated by subtracting the rate of mitochondrial matches to the consensus sequence from 1 (smooth bars) and based the heterozygosity of the X-chromosome of male individuals (textured bars). Error bars indicate the 95% confidence interval.

Out of 10 individuals included in this study, both the cochlea and ossicles produced enough data to call mitochondrial DNA (mtDNA) haplogroups, assess damage patterns at the terminal nucleotide of the molecule, and make contamination estimations for seven individuals; these individuals are henceforth referred to as the ‘working individuals.’ One individual from Thailand produced marginal data that allowed the same analyses, but produced a larger error interval for the mtDNA contamination estimate – calculated as 1 minus the rate of mitochondrial matches to the consensus sequence (Fu et al. 2013) – only when the ossicles were used; two individuals, both from Yemen, did not produce enough data to allow for the determination of the mtDNA haplogroup or contamination estimates. Both the cochlea and ossicles were therefore considered to have ‘failed’ our analysis for these latter individuals. We performed Wilcoxon Signed-Rank tests to compare the data generated using the ossicles and cochlear samples.

We obtained an average endogenous DNA yield of 45.87% for the seven working cochlea samples and 51.30% for the corresponding ossicles (Table 1, Figure 2 Panel A) (p=0.2969 for the difference; Supplementary Table 3). Complexity, defined here as the percentage of unique reads expected out after down-sampling to 500,000 sequences that align to the ∼1.2 million targeted SNPs, is a potentially more informative metric for comparing performance between the cochlea and ossicles because it is directly related to the maximum amount of sequencing data the extract or library can possibly yield and is not biased by differences in sequencing depth across samples. The average complexity for cochlea and ossicles was 87.1% and 86.0%, respectively (Table 1, Figure 2 Panel B); this difference is also non-significant (p=0.4688; Supplementary Table 3). Overall, these results suggest that the data generated using ossicles is comparable to that generated using the cochlea. Any minor differences are likely due to chance rather than a systematic difference in DNA preservation between the cochlea and ossicles.

The average mtDNA coverage was 525× for the seven working petrous samples and 486× for the corresponding ossicles (Supplementary Table 2), which were not significantly different (p=0.6875; Supplementary Table 3). The average coverage of the ∼1.2 million targeted SNPs from across the genome was 1.53× for the seven working petrous samples, and 1.47× for the ossicles (Table 1); on average, 727,500 SNPs were called when the cochlea was used and 714,312 were called when the ossicles were used (Table 1). Both of these differences were non-significant (p=0.9375 and 0.6875, respectively; Supplementary Table 3).

For a sample from burial phase Middle Period VII at Ban Chiang, northeast Thailand (BCB 26), the cochlea failed to produce enough data even for estimating contamination, with only 266 nuclear SNPs covered; however, we observe a ∼46-fold increase in SNPs hit associated with the use of the ossicles (12,438 SNPs) (Table 1). In addition, the mitochondrial coverage was seen to increase from 0.08x with the cochlea to 5.15x with the ossicles, an increase of ∼63-fold (Table 1, Supplementary Table 2). Looking further into this data increase, we note a ∼4-fold decrease in frequency of deamination at the terminal base (from 6.40% to 1.30%) for the nuclear data as well as a high mitochondrial contamination estimate (point estimate, 6.0%; 95% confidence interval: 2.8–12.3%), which may indicate the presence of DNA contamination (Table 1, Figure 2). Because of this, we are unable to equate the increase in data to the use of the ossicle.

For the seven working samples, the average deamination frequency was slightly reduced from 12.32% to 11.28% when the ossicles were used, a decrease (Table 1, Figure 2 Panel C) that, although small, was statistically significant (p=0.0313; Supplementary Table 3). Mitochondrial contamination estimates (inferred by identifying mismatches to the mtDNA consensus sequence (Fu et al. 2013)) increased from an average of 0.63% to 1.44%, (Table 1, Figure 2 Panel D) with a significant p-value of 0.0469 (Supplementary Table 3). This change was driven by a single individual (818), which exhibited increased contamination in the ossicle relative to the cochlea (Table 1, Figure 2, Supplementary Table 3). Contamination based on the heterozygosity rate of the X-chromosome (a test only applicable to males) (Korneliussen et al. 2014) averaged 0.52% for the cochlea and 1.12% for the ossicles (or excluding individual 818, 0.53% and 0.40%, respectively), a non-significant change (p=0.625 for the full test and 0.125 without individual 818) (Table 1, Figure 2, Supplementary Table 3). The overall low levels of contamination are also supported by consistency in the estimation of mtDNA haplogroups and molecular sex for all cochlea-ossicles pairs (Table 1, Figure 2, Supplementary Table 2).

## DISCUSSION

### DNA recovery from the auditory ossicles

This study presents a direct comparison of DNA recovery from the ossicles and corresponding cochlear bone using archaeological specimens that originate from varying geographic and temporal contexts and offers several new insights. First, we demonstrate that the ossicles perform comparably to the cochlea in terms of ancient DNA recovery regardless of sample preservation. Focusing on seven individuals from whom we were able to generate enough working ancient DNA data to call mtDNA haplogroups, assess damage pattern, and make contamination estimates, we find that the use of the cochlea or ossicles from each individual produces similar amounts of endogenous DNA, mtDNA coverage, nuclear SNP coverage, and number of SNPs called. We demonstrate that there is no substantial reduction in data quantity or complexity associated with the analysis of the ossicles instead of the cochlea. Second, although we find that the ossicles show a slight reduction in the frequency of deamination (a signal of ancient DNA authenticity) compared to the corresponding cochlea, the amounts of contamination estimated using both mtDNA and heterozygosity on the X chromosome are comparable. Considered together, our data suggest that there is little reduction in data quality associated with the analysis of the ossicles instead of the cochlea. We conclude that the auditory ossicles, when present, are an alternative optimal skeletal element that can be used in ancient DNA research in place of the cochlea

Though they are small, often isolated, and can be accessed without significant impact to larger, morphologically-informative parts of the skeleton, the use of ossicles for ancient DNA analysis still requires the destruction of human skeletal material that may be anthropologically valuable. Ossicles have previously been used in studies of comparative morphology; most notably, they have provided insight into morphological differences and functional similarities in the middle ear of Neandertals and anatomically modern humans, which has implications for understanding the auditory capacity of extinct hominins (e.g., Stoessel et al. 2016). For this reason, we encourage all researchers contemplating ancient DNA analysis to balance their analytical goals with the impact that sampling for ancient DNA analysis will have on future availability of material.

In light of these findings, we suggest that archaeologists and curators attempt to identify and preserve auditory ossicles whenever possible. Ideally, ossicles would be identified and collected during archaeological recovery of human skeletal remains in a way that minimizes the introduction of contamination. This includes wearing disposable medical gloves that are changed frequently when handling samples, avoiding washing skeletal material with water, and storing samples in a cold, dry place as soon as possible (Llamas et al. 2017).

The use of ossicles for ancient DNA analysis will contribute to the successful analysis of skeletal material that does not have a petrous bone present, or sets of remains that have a petrous bone that cannot be processed in a destructive manner for ancient DNA research (for example, those that may be morphologically-intact and displayed in museum collections). On a broader level, the identification of the ossicles as an alternative optimal skeletal element for ancient DNA analysis contributes to the reduction in the amount of damage inflicted to human skeletal samples for the purposes of ancient DNA analysis. It is another step toward the preservation of DNA-rich and anthropologically-valuable skeletal material for future studies that may benefit from methodological improvements that are unknown at present.

## METHODS

### Sample Selection and Preparation

The number of ossicles collected for each of the 10 archaeological samples varied (see Table 1), but the incus and malleus were identified and collected most frequently (n=10 and n=8, respectively) while the stapes was identified and collected least frequently (n=2), likely due to its diminutive size and fragility. In most cases, we recovered the ossicles while following the standard cochlea sampling procedure (Pinhasi et al. 2019). In other cases, we intentionally dislodged the ossicles from the skull for the purpose of this study; in most of these instances, the ossicles were partially visible within the external auditory meatus. To dislodge the ossicles, we cleaned a small engraving burr (described in Sirak et al. 2017) by wiping it with a diluted bleach solution (∼10% concentration). We placed the cleaned burr inside the external auditory meatus and gently manipulated it within the inner ear canal. This caused no apparent damage to the ossicles or to the cranium from which they were retrieved. All ossicles were immediately placed into a sterile 2.0mL tube upon their removal from the ear canal.

The preparation of all skeletal material for ancient DNA analysis was carried out in dedicated cleanrooms at University College Dublin (UCD) or at the University of Vienna following standard anti-contamination protocols (e.g., Hofreiter et al. 2001; Poinar 2003; Llamas et al. 2017). All petrous bones were processed following a standard protocol (Pinhasi et al. 2019). This protocol uses a dental sandblaster to systematically locate, isolate, and clean the cochlea, which is then milled to homogeneous bone powder. Approximately 50 mg of bone powder from the cochlea (range: 47–56 mg) was aliquoted for DNA extraction. Complete auditory ossicles were decontaminated through exposure to UV irradiation for 10 minutes on each side; after noting a substantial reduction in amount of bone powder associated with the milling of complete ossicles to bone powder during pilot work, we chose not to grind the ossicles to a fine powder, instead placing them inside a new sterile 2.0mL tube following decontamination with UV irradiation. The tubes that included the whole ossicles or petrous bone powder were then taken to a separate ancient DNA clean room for DNA extraction and preparation of sequencing libraries.

### DNA Extraction

DNA was extracted from the cochlear bone powder and the whole auditory ossicles in ancient DNA facilities at the University of Vienna following a standard ancient DNA extravtion protocol (Dabney et al. 2013) with a modification (Korlević et al. 2015) that uses the tube assemblies from the High Pure Viral Nucleic Acid Large Volume kit (Roche, 05114403001). The intact ossicles were placed in the extraction buffer, and completely dissolved during the incubation period in most cases. Lysates were washed twice with 650 µL of PE buffer (Qiagen) and spun through the columns at 6000 rpm for 1 minute. After being put in a fresh 1.5mL collection tube, 25µL of TET buffer was pipetted on the dry spun MinElute columns’ silica membrane. After a 10-minute incubation at room-temperature, the columns were spun at maximum speed for 1 minute. The elution step was repeated to give a final volume of 50µL of DNA extract. A negative control that contained no bone material was included with each extraction batch.

### Library Preparation

Next generation sequencing libraries were prepared in ancient DNA facilities at Harvard Medical School from all extracts and controls using a library preparation method optimized for ancient DNA (Rohland et al. 2015). This protocol uses a partial-UDG treatment that causes characteristic C-to-T ancient DNA damage to be restricted to the terminal molecules while nearly eliminating it in the interior of the DNA molecules so that the library can be used to test for ancient DNA authenticity. 10μL of DNA extract was used as input during library preparation. Libraries were enriched for ∼1.2 million nuclear sites across the genome (‘1240K capture’) in addition to sites on the human mitochondrial genome (Fu et al. 2013, 2015; Haak et al. 2015; Mathieson et al. 2015). Enriched libraries were sequenced on an Illumina NextSeq500 instrument, with 2×76 cycles and an additional 2×7 cycles used for identification of indices. In addition, a small proportion of reads were generated from unenriched versions of each library. This unenriched (‘shotgun’) data was used to estimate the proportion of endogenous molecules in each library.

### Data Processing

Following sequencing, we trimmed molecular adapters and barcodes from sequenced reads prior to merging forward and reverse reads using custom software (https://github.com/DReichLab/ADNA-Tools). We allowed up to three mismatches of low base quality (<20) and up to one mismatch at higher base quality (≥20), ensuring that the highest base quality in the overlap region was regained. We aligned reads to the mitochondrial *RSRS* genome (Behar et al. 2012) and to the *hg19* human reference sequence with the *samse* command in *bwa* (v0.6.1) (Li and Durbin 2009).

We used the tool ContamMix (Fu et al. 2014) to determine the rate of matching between the consensus RSRS sequence and reads which aligned to the mitochondrial genome. We determined the rate of C-to-T substitution at the terminal ends of each molecule using PMDtools (https://github.com/pontussk/PMDtools; Skoglund et al. 2014). We used the tool ANGSD (Korneliussen et al. 2014) to determine the amount of contamination in the X-chromosome of individuals identified as genetically male. The complexity of the sample was assessed by quantifying the number of unique reads expected from a pre-determined number of reads hitting target.

## DATA ACCESS

Data are available at the European Nucleotide Archive under accession number PRJEB32751.

## ACKNOWLEDGEMENTS

This work was partially supported by a European Research Council starting grant ADNABIOARC 263441 to R.P, a National Science Foundation (NSF) Doctoral Dissertation Research Improvement Grant BCS-1613577 to K.Si, an Irish Research Council grant GOIPG/2013/36 to D.F., a graduate student fellowship from the Max Planck-Harvard Research Center for the Archaeoscience of the Ancient Mediterranean (MHAAM) to E.H, and Russian Foundation for Basic Research grants 18-00-00360, 18-09-00349 to V.M. T.H. and T.S. were supported by grants from the Hungarian Research, Development and Innovation Office, project numbers FK128013 and TÉT_16-1-2016-0020. D.R. is an investigator of the Howard Hughes Medical Institute. The authors would like to thank Canterbury Archaeological Trust for permission to analyze sample 1257 (I4491); full details about this sample can be found in the CAT PXA report (Helm et al. 2017).

## SUPPLEMENTARY MATERIAL

### Pilot Work

Five archaeological samples representing a range of geographic locations were selected for a pilot project aimed at obtaining initial insight into use of the ossicles for ancient DNA analysis. We chose samples based on their age and depositional contexts to represent a range of molecular preservation (sample information provided in Supplementary Table 1). All specimens had at least two ossicles, and one petrous pyramid from the same individual was selected for comparative analysis. Skeletal material was processed in dedicated ancient DNA clean rooms at University College Dublin following standard anti-contamination protocols (Hofreiter et al. 2001; Poinar 2003; Llamas et al. 2017). Petrous bones were processed as described in Pinhasi et al. (2019) to create bone powder, and complete auditory ossicles were decontaminated through exposure to UV irradiation for 10 minutes on each side and milled to fine powder. DNA extraction and library preparation followed standard ancient DNA protocols, described in the following section. All extraction and library preparation took place in a separate clean room from that used for processing bones and also followed standard anti-contamination protocols.

We generated raw sequencing data for this pilot work using low-coverage whole-genome shotgun sequencing on the Illumina MiSeq and NextSeq platforms. Data were processed using a custom bioinformatics pipeline to enable a basic comparison of endogenous DNA yield from the cochlea and from the auditory ossicles (Supplementary Table 1). Our results suggested that the auditory ossicles were approximately equivalent to the cochlea for endogenous DNA preservation, with the difference in endogenous DNA content ranging between a 0.17-fold decrease and a 0.3-fold increase (Supplementary Table 1). The endogenous DNA yields ranged from 0.16 to 68.19%, with a median of 54.68%, and no substantial difference between the ossicles and cochlea detected (Supplementary Table 1). We identified damage patterns consistent with expectations for ancient DNA in the sequencing data generated using both the ossicle and cochlea samples, with an average substitution frequency on the 5’-end of the DNA molecule of 14.50% for the ossicle samples and 14.40% for the petrous bone samples (Supplementary Table 1). Like endogenous yield, this difference is not substantial. Overall similarity in endogenous yield and damage frequencies between the auditory ossicle and cochlea samples from the same individual supported our hypothesis that auditory ossicles may also be an effective substrate for ancient DNA analysis.

